# Topic modeling for multi-omic integration in the human gut microbiome and implications for Autism

**DOI:** 10.1101/2022.09.30.509056

**Authors:** Christine Tataru, Marie Peras, Erica Rutherford, Kaiti Dunlap, Xiaochen Yin, Brianna S. Chrisman, Todd Z. DeSantis, Dennis P. Wall, Shoko Iwai, Maude M. David

## Abstract

While healthy gut microbiomes are critical to human health, pertinent microbial processes remain largely undefined, partially due to differential bias among profiling techniques. By simultaneously integrating multiple profiling methods, multi-omic analysis can define generalizable microbial pro-cesses, and is especially useful in understanding complex conditions such as Autism. Challenges with integrating heterogeneous data produced by multiple profiling methods can be overcome using Latent Dirichlet Allocation (LDA), a promising natural language processing technique that identifies topics in heterogeneous documents.

In this study, we apply LDA to multi-omic microbial data (16S rRNA amplicon, shotgun metagenomic, shotgun metatranscriptomic, and untargeted metabolomic profiling) from the stool of 81 children with and without Autism. We identify topics, or microbial processes, that summarize complex phenomena occurring within gut microbial communities. We then subset stool samples by topic distribution, and identify metabolites, specifically neurotransmitter precursors and fatty acid derivatives, that differ significantly between children with and without Autism. We identify clusters of topics, deemed “cross-omic topics”, which we hypothesize are representative of generalizable microbial processes observable regardless of profiling method. Interpreting topics, we find each represents a particular diet, and we heuristically label each cross-omic topic as: healthy/general function, age-associated function, transcriptional regulation, and opportunistic pathogenesis.

## Introduction

Autism is a complex neurodevelopmental condition that occurs in 1 out of every 54 children in the United States. It is characterized by a specific set of behaviors, although many autistic people may exhibit only a subset of them ***Lord et al. (1989)***. Autism is frequently called “Autism Spectrum Disorder” in the scientific literature, however, surveys and public opinion from people in the community have demonstrated that the preferred term is simply “Autism” ***Organization for Autism Research (2020)***.

Autistic people have a statistically higher likelihood to experience gastrointestinal (GI) issues ***Wasilewska and Klukowski (2015); Kohane et al. (2012)***. These GI issues can include symptoms such as chronic constipation, diarrhea, abdominal pain, and potential signs of GI inflammation such as vomiting and bloody stools ***Hsiao (2014)***. There is also growing evidence to suggest a link between gut microbiome dysbiosis and Autism. Fecal microbiota transplants in children with Autism have demonstrated some improvements in GI symptoms and a shift in Autism-associated behaviors ***Kang et al. (2017, 2019)***. Probiotic supplementation has also been observed to improve GI symptoms as well as multisensory processing and adaptive functioning in autistic preschoolers ***Santocchi et al. (2020)***. However, the specifics of this relationship are still little understood. Many studies using 16S sequencing to profile microbial communities have reported differences in the abundance of certain species and genera - often these reports are not reproducible in independent studies ***Nitschke et al. (2020); Iglesias-Vázquez et al. (2020); Xu et al. (2019)***.

Molecular-level processes have also been implicated in Autism. Neurotransmitter biosynthesis and breakdown, specifically tryptophan and glutamine metabolism, are often observed to be dysregulated in Autism and potentially influenced by shifts in the gut microbiome ***Al-Otaish et al. (2018), Marotta et al. (2020), Cochran et al. (2015), Horder et al. (2013), Strandwitz et al. (2019), Yano et al. (2015)***. Additionally, the chance of mitochondrial disease within the autistic population is about 5.0%, 500 times higher than that found in the general population (0.01%), and 30% of children with Autism may experience metabolic abnormalities ***Cheng et al. (2017)***. The metabolisms of the resident gut bacteria is suggested to be involved in both of these phenomena respectively ***Hu et al. (2020)***. Our understanding is hindered by the heterogeneity of the categorization of Autism, the human gut microbial ecosystem, and the multiple profiling techniques that may be used to measure microbiomes.

There are certain techniques in computer science, specifically in natural language processing, that excel at summarizing highly heterogeneous data. The topic modeling strategy used here, Latent Dirichlet Allocation (LDA), was originally used to identify topics within heterogeneous written text documents ***Blei et al. (2003)***. Based on the shared distribution of words across documents, LDA defines a pre-determined number of topics, each of which is a probability distribution across words. Documents are simultaneously defined by a probability distribution across topics. In this way, a document from a food blog may be 60% about restaurants, and 40% about travel. This mixed membership model set up allows models to capture complex phenomenon as would be expected in a microbial ecosystem.

In this study, we pursued two objectives. First, we sought to deepen our understanding of related variables within the human gut microbiome as represented by multiple omic profiling technologies. We define an “omic” as one sample by feature table as produced by one profiling method (i.e. 16S rRNA, shotgun metagenomics, shotgun metatranscriptomics, and untargeted metabolomics). To address this objective, we applied Latent Dirichlet Allocation, a mixed membership statistical method, defining microbial features (i.e. amplicon sequencing variants (ASVs), Kegg Orthologs (KOs), or compounds) as words, and samples as documents. This topic analysis defined specific sub-processes or topics that were represented in a robust manner across multiple gut mi-crobiome profiling techniques. Second, we evaluated potential associations between features of the gut microbial ecosystem and Autism. To do this, we first used the identified microbiome topics to cluster samples with similar”microbial landscapes” together, then applied traditional differential abundance analysis techniques.

Our first objective was motivated by the fact that human gut microbiomes are highly heteroge-neous within and between individuals on a taxonomic and functional level. Clustering approaches are commonly used to define community structure and have led to the adoption of the concept of enterotypes, distinct sub-types of microbiomes that are dominated by either Bacteroides, Pre-votella, or Ruminococus genera ***Arumugam et al. (2011)***. However, while this categorization serves as an important dimensionality reduction tool for these complex datasets, it tends to oversimplify the community structure, and hides complexity in the microbial communities that exist in the space between enterotypes ***Knights et al. (2014)***. Moreover, while clustering approaches can identify frequently co-occurring species, they do not identify species that share similar co-occurrence patterns but do not themselves directly co-occur (as might be seen in species with similar function) ***Symul et al. (2022)***. Newer work argues that host-associated microbiomes should be considered to be on a spectrum, and that mixed membership models, and in particular topic models, are powerful and robust tools to learn and define that spectrum ***Knights et al. (2014); Sankaran and Holmes (2018); Deek and Li (2021); Breuninger et al. (2021); Sommeria-Klein et al. (2020); Okui (2020)***.

Our second objective, to identify associations between Autism and gut microbial features, was motivated by the high heterogeneity between individual microbiomes, and was enabled by topic modeling. Reports of how the gut microbiome relates to Autism are highly variable across the literature, and while many agree that the gut-brain axis plays a role, there is only moderate consensus about the specific bacteria and processes involved ***Nitschke et al. (2020); Iglesias-Vázquez et al. (2020); Xu et al. (2019)***. Differential abundance analysis between highly heterogeneous samples provides limited power and is more likely to produce spurious positive results. To address this, we use topics identified in multi-omic data to cluster samples, and perform differential abundance analysis on each cluster independently, making each group less heterogeneous in microbial structure.

## Results

### Topic modeling for multi-omic integration

We used Latent Dirichlet Allocation (LDA), a form of unsupervised topic modeling, as an integrative approach to reduce the thousands of features across 16S rRNA(16S), shotgun metagenomic(MTG), shotgun metatranscriptomic (MTT), and untargeted metabolomic (MBX) profiling datasets all performed on the same stool samples from children with and without Autism. LDA was originally used for topic modeling in natural language processing where topics are defined by distributions across a vocabulary and documents are modeled as deriving from a distribution of topics. In this study case, we treated samples as documents and omic features as words, and we used the counts of features across samples to learn topics per omic (Figure 1 A). After model fitting, each topic is a weighted vector of feature count probabilities, and each sample is a weighted vector of topic probabilities. Each sample has 24 total associated vectors: 4 original omic count vectors, 4 16s topic vectors, 5 MTG topic vectors, 4 MTT topic vectors, and 7 MBX topic vectors (Figure 1 B). The number of topics were selected to minimize differences between true and model-simulated data (see Methods). Remarkably, topics learned from different omics correlated significantly with one another in true data, but not in data simulated using null models (see Methods), allowing us to define cross-omic topics, or sub-processes that are observable at multiple levels of measurements (Figure 1 C, Supp. Figure 1). Using the topics as representative of broad microbial processes, we split samples into two main groups based on their topic distribution (Figure 1 D). Samples in the same topic-derived clusters shared similar dietary patterns (Supp. Figure 2). Clustering samples using topics highlighted two distinct Autism-related metabolic profiles: sample cluster 1, which associated with healthier eating habits, implicated neurotransmitter precursors and sample cluster 2, which associated with less healthy eating habits, implicated fatty acids and their derivatives. Lastly, topic interpretation revealed four main processes reflected in all omics: healthy/general function, age-associated function, transcriptional regulation, and opportunistic pathogenesis.

**Figure 1.**
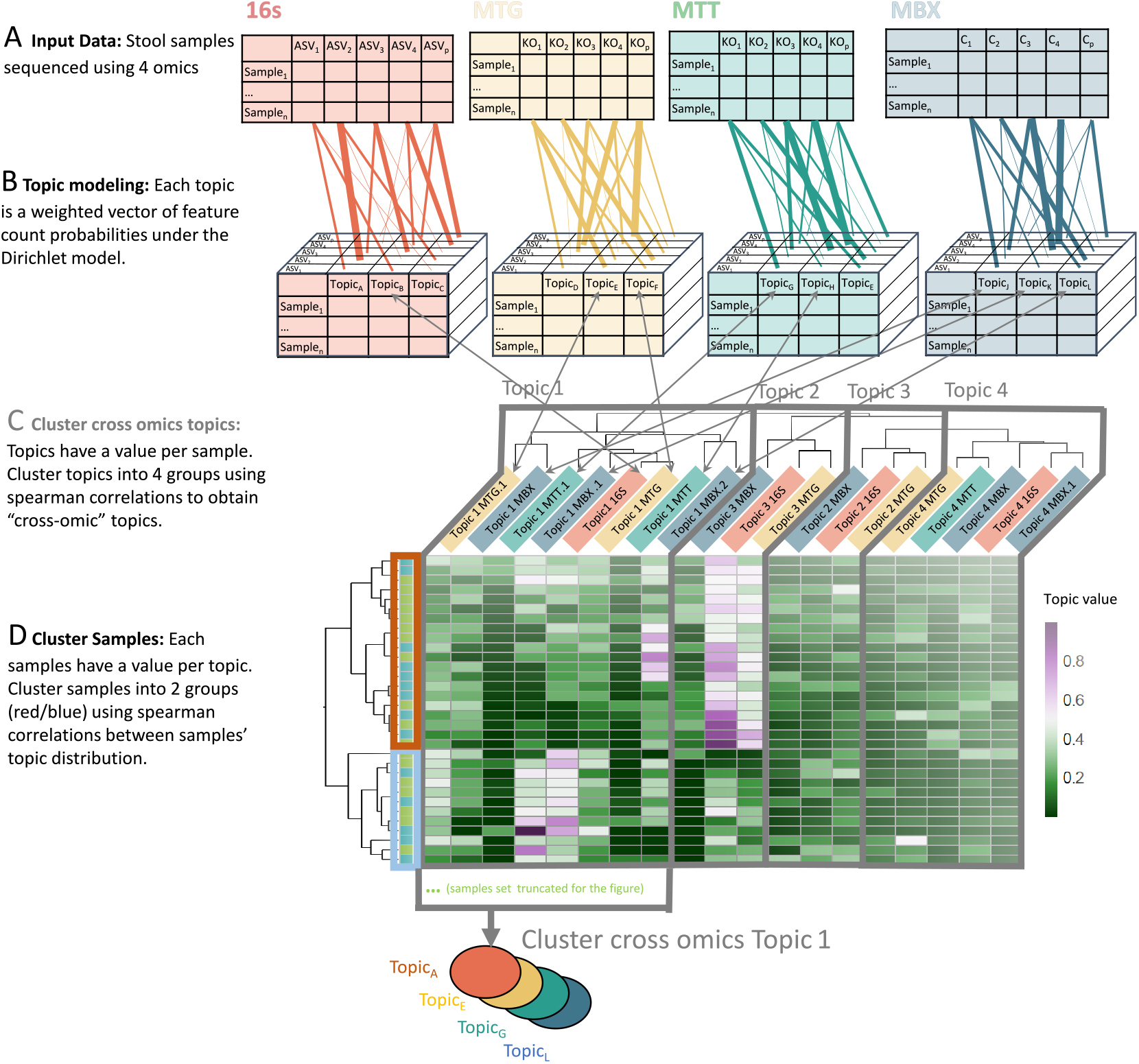
Topic modeling process. Stool samples were measured using 4 techniques, 16s, metagenomics (MTG), metatranscriptomics(MTT), and metabolomics (MBX) (A). Latent Dirichlet Allocation (LDA) was applied to each dataset independently to model latent variable “topics” from count data. Each sample is modeled as a Dirichlet distribution of topics, and each topic is modeled as a Dirichlet distribution of features (ASV, KO, or C) (B). Topics from each omic were then clustered by their distribution across samples using hierarchical clustering on spearman correlations (C). We call the cluster of topics “cross-omic topics”, and discuss their interpretations and biological implications. Multiple topics from an omic may be included in the same cross-omic topic, denoted as “.1” or “.2” etc. Samples were clustered using hierarchical clustering on spearman correlations between their topic distributions (D). Groups enabled comparisons between samples with similar diets and similar ‘microbial feature landscapes’.

### Modeling identifies cross-omic topics that correspond to diet and microbial functions

We interpreted topics by observing which features have highest attribution weights for each topic,and also by correlating topic distributions across samples to dietary characteristics (Figure 2). We find that dietary variables cluster into two main groups (Figure 2E) with the consumption of fruits, vegetables, fermented vegetables, seafood, meat, home-prepared meals, probiotics, vitamin B, and vitamin D constituting one group, and consumption of sugary food, sweetened drinks, starchy food, dairy, whole-grains, pre-packaged meals, and restaurant food constituting another. It should be noted that dietary variables are not included into the topic modeling process itself to focus analysis on the true microbial landscapes and erase bias due to self-reported data and pre-conceived notions of healthy eating.

**Figure 2.**
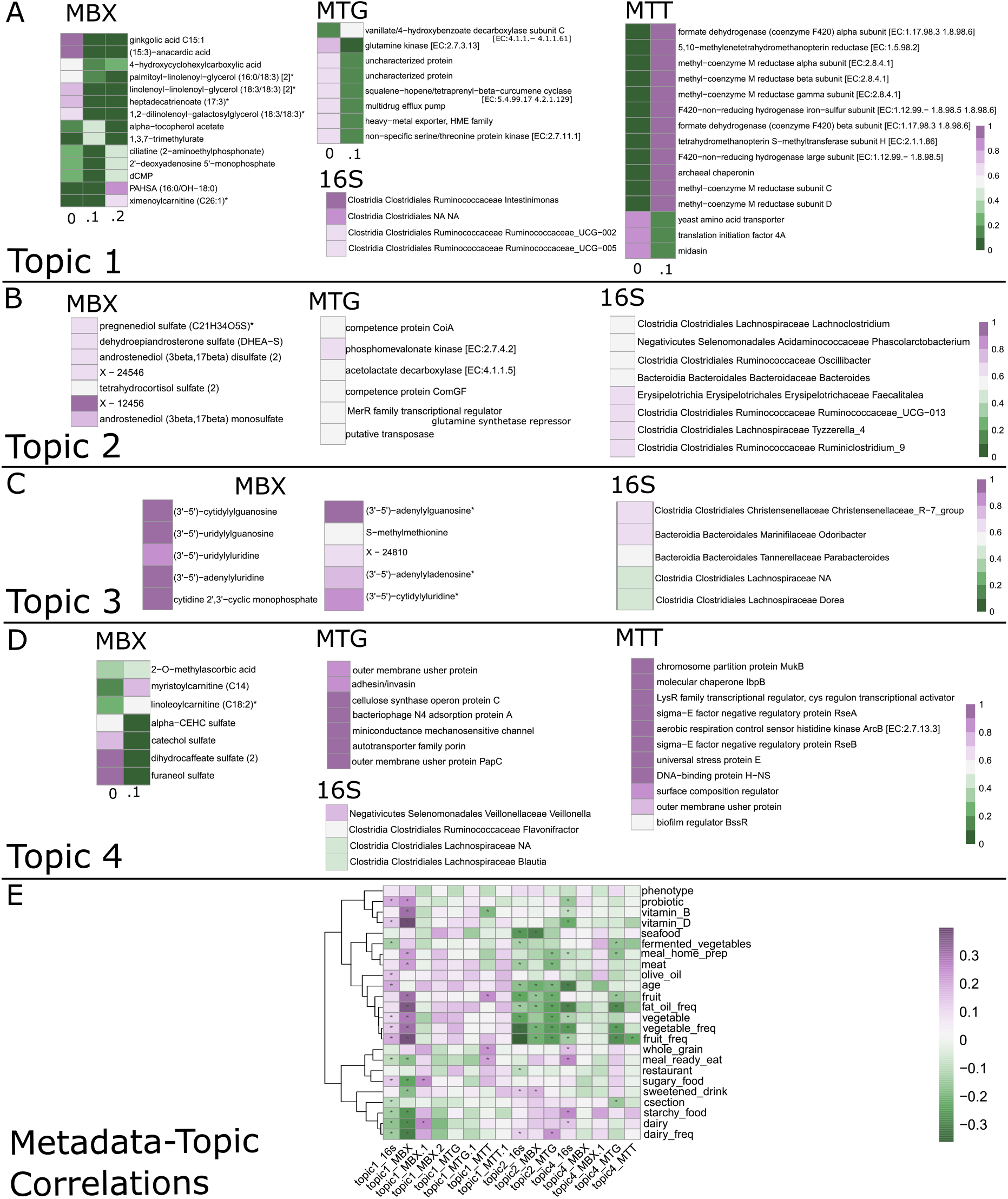
Topic interpretation using feature weights. Each topic may be defined by those features with the highest weights. Features detailed here had weights with max value more than 1.5 standard deviations over the median (exceptionally high value) and high values along axes of maximal variation between topics (exceptionally attributable to a single topic). Topic interpretations are for MBX (A), 16S (B), MTT (C) and MTG (D). Topic correlations with dietary features are in (E). Dietary variables with keyword “freq” represent long-term self-reported variables, while variables without “freq” are about the most recent week before sampling. Cells are colored by topic value (probability between 0 and 1) and a star signifies a significant (p<0.05) spearman correlation

Topic1 is characterized by high consumption of the first, healthier, group of dietary variables (Figure 2E), as well as with bacterial alpha diversity (Supp. Figure 3). There are three MBX topics that cluster into cross-omic Topic1. They are characterized by high values of fats, specifically 18:3 containing glycerols, anacardic and heptadecatrienoic acids, PAHSA, ximenoylcarnitine, as well as alpha-tocopherol acetate, trimethylurate, and AAMU (5-acetyleamino-6-amino-3-methyluracil) (Figure 2). There are two MTG topics that cluster into cross-omic Topic1. They are characterized by ubiquitous enzymes like glutamine and serine/threonine kinases and heavy metal and drug exporter pumps. There are two MTT topics that cluster into cross-omic Topic1 as well. They are characterized by high values of methyl-coenzyme M subunits used in methane metabolisms, as well as functions like yeast plasma membrane ATPase, yeast amino acid transporter, eukaryotic translation initiation factors, pyruvate decarboxylase, heat shock protein, and MFS sugar transporters. There is one topic from 16s that clusters into cross-omic Topic1; it is high in values for species from Ruminococcacea, as well as other unidentified Clostridiales order members.

Additionally, we found Topic1 values to be correlated with specific behavioral characteristics from the parent-reported behavioral questionnaire (see Methods). In particular, Topic1 values correlated with observed imaginative play, both along and with others, as well as language and speech skills and a lack of self harm behavior (Supp. Figure 4). It should be noted that these behaviors and topics may be related to age, although significance of correlations between Topic1 and age vary (Supp. Figure 5).

Cross-omic Topic2 is characterized by high consumption of the second, less healthy, group of dietary variables (Figure 2 E). In MBX data, we see high values of steroid sulfates, specifically an-drostenediol mono and di sulfate, tetrahydrocortisol sulfate, pregnenediol sulfate, and DHEA-S (Figure 2 B). In MTG, we see proteins used by bacteria for horizontal gene transfer, namely competence proteins and transposases, as well as a glutamine synthetase repressor, acetolactate decarboxylase, and phosphomevalonate kinase. In 16s, we see high Ruminiclostridium species, Blautia and Tyzzerella and Faecalitalea genera members. Topic2 is not represented in MTT data. Topic2 is also the only cross-omic topic consistently correlated (negatively) with age (Supp. Figure 5)

Topic3 is weakly associated with the first, healthy, set of dietary variables (Figure 2 E). In MBX data, we see high values of dinucleoside monophosphates (Figure 2 C). In MTG data, we see higher values of sensor histidine kinase DegS and manganese/zinc/iron transport permease protein. In MTT data, Topic3 is not clearly defined by any set of features, but is dramatically the topic assigned the highest attribution across all samples (the chance of a sample having high Topic3 values is very high) (Supp. Figure 6). In 16s data, we see high values for two species, one from the family Christensenellaceae and the other from Odoribacter.

Topic4 is characterized less strongly by metadata variables, but still represents lower intake of vegetable, fruit, fat/oil, meat, and home prepared meals (Figure 2 E). In MBX data, we see high values of sulfates, specifically catechol, dihydrocaffeate, and furaneol sulfates (Figure 2 D). Additionally, myristoylcarnitine, linoleoylcarnitine, 7-ketolithocholate, 3-hydroxyisobutyrate, and 2-O-methylascorbic acid are all high in Topic4 MBX. In MTG, we see a strong contribution from genes associated with pathogenesis, specifically autotransporter family proteins, usher proteins, adhesin/invasin, and bacteriophage proteins. In MTT, we see similarly high values of universal stress protein E, outer membrane usher protein, biofilm regulator BssR, as well as regulatory proteins RseB, molecular chaperone IbpA, and chromosome partition protein MukB. Lastly, 16s data shows particularly high values for a Veillonella species, as well as for species from the Blautia, Faecalibacterium, Intestinibacter, and Anaerostipes genera.

### Subsetting population by topics reveals metabolic differentials between Autism and Typically Developing cohorts

We defined two major clusters of samples using hierarchical clustering oftopic distributions (Figure 3 A). We found that 92% of families (88 families) clustered into the same groups as each other, while only 8% (8 families) clustered separately (data not shown). Before clustering, autistic and typically developing participants had large differences in multiple dietary variables that significantly affected multi-omic features (Supp. Figure 2). After separating samples into subpopulations, we found that metadata differences between autistic and typically developing phenotypes became less stark (Supp. Figure 7). The exception to this statement was autistic individuals in Cluster2 still ate less dairy than their typcally developing counterparts. This grouping therefore allowed us to account for differences in dietary patterns in our analysis.

**Figure 3.**
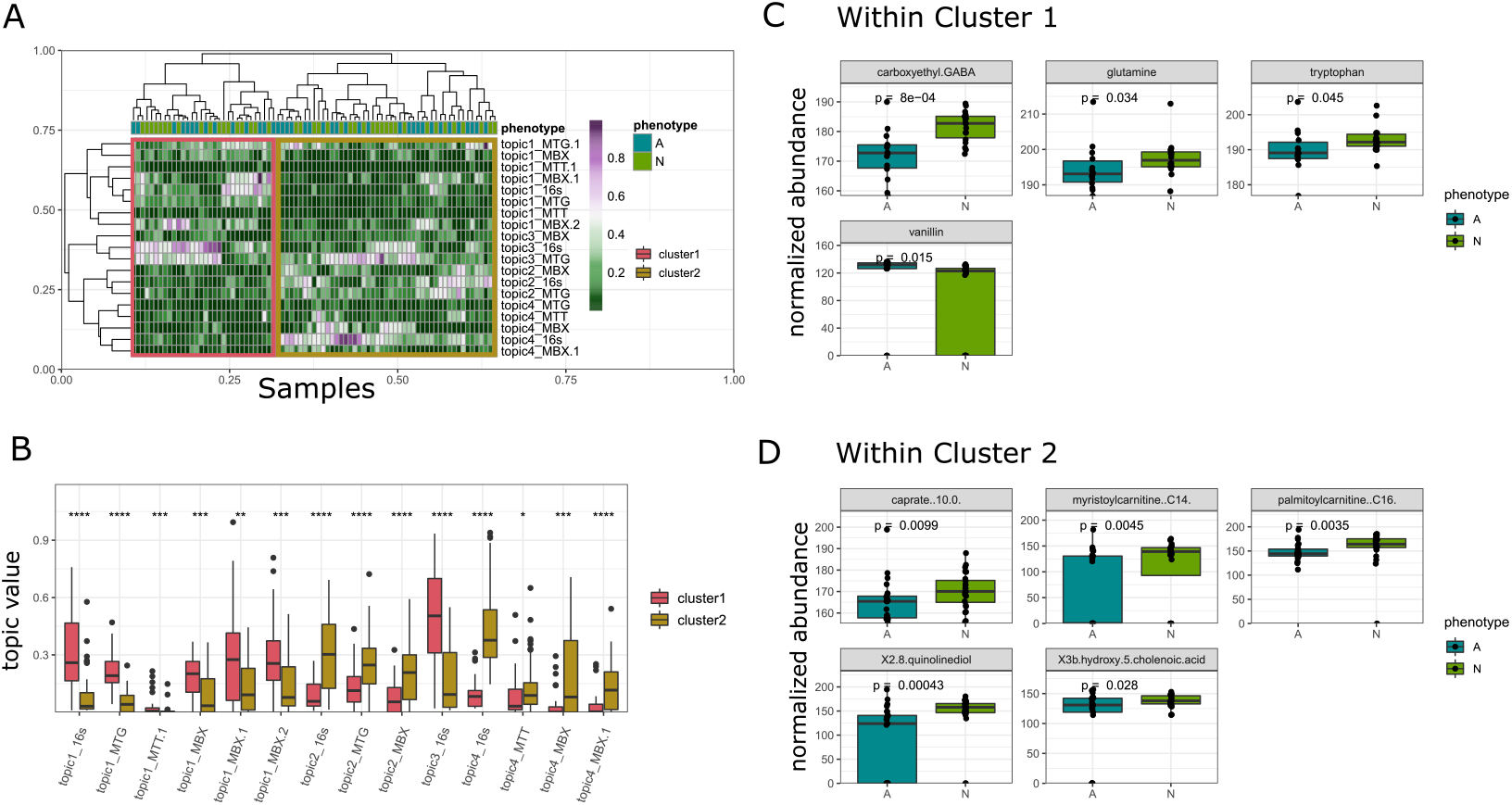
Differential analysis of metabolites between autistic and typically developing children within sample clusters as determined by topics. Hierarchical clustering on samples’ topic distribution produces two clusters (A). Cluster1 is defined by high topics 1 and 3, while cluster 2 is defined by high topics 2 and 4 (B). In cluster 1, CEGABA, glutamine, tryptophan, and vanillin are differentially abundant (C). In cluster 2, caprate, myristoylcarnintine, palmitoylcarnitine, quinolinediol, and 3hydroxy5cholenoic acid are differentially abundant (D). Features are first filtered, and then p values are reported using Wilcox rank sum test. Multiple single-omic topics may belong to the same cross-omic topic, denoted here as “.1” or “.2”

We find samples in cluster1 have high cross-omic topic1 values, while samples in cluster2 have high cross-omic topic2 and topic4 values (Figure 3 B). Differential analysis between autistic and typically developing individuals within these subset populations reveals distinct metabolic signatures. In Cluster1, autistic individuals have significantly lower stool abundances of neurotransmitter precursors, namely caroxyethyl GABA, glutamine, and tryptophan, and higher abundance of vanillin (Figure 3 C). In Cluster2, autistic individuals have significantly lower stool abundances of fatty acids (caprate), acylcarnitines (myristoylcarnitine, palmitoylcarnitine), quinolinediol, and 3-hydroxy-5-cholenoic acid (Figure 3 D).

Additionally, we observed differentials in metagenome, metatranscriptome, and 16s features between autistic and typically developing individuals in each subgroup (Supp. Figure 8)).

## Discussion

The gut microbiome has been implicated in a plethora of complex and idiopathic conditions, but thus far remains challenging to interpret due to the high variability and many avenues for analysis. We may observe microbiome structure and function using 16s rRNA amplicon sequencing, shotgun metagenomics, shotgun metatranscriptomics, and untargeted metabolomic profiling (among many others), and each of these data modalities reveals a different understanding of the composition and dynamics of the microbial community at large. While none of these methods singularly assures accurate conclusions, this paper simultaneously integrates all four omic data modalities to create a more complete and unbiased representation of microbiome structure and function. Our approach reveals which dietary habits are reflected across the microbiome to what extent, and presents an objective approach to subtyping a population based on microbiome composition and function.

We define ‘cross-omic topics’ that summarize patterns observed in 16s, metagenome, metatranscriptome, and metabolomic data, and their relationship with each other. We find that topics correlate with dietary variables, and these correlations can be used to group dietary habits in a way that reflects current dietary health standards. We associate features from each omic with those groups, and hypothesize that these feature groups may be used to obtain a more complete and data-driven understanding of overall health of the microbiome, especially in complex diseases as-sociated with microbiome dysbiosis. We then clustered stool from autistic/typically developing participants using topic values, and found two distinct subpopulations. Metabolic differences between autistic and typically developing individuals within each subpopulation implicated neurotransmitter precursosr and medium to long chain fatty acid availability as relevant factors in Autism worthy of further exploration. We argue that this type of topic modeling approach could be used in other complex disease studies to integrate multi-omic data and more accurately elucidate the relevant factors in the human-microbiota relationship.

### Value of LDA

Latent Dirichlet Allocation (LDA) has been used with great success to identify topics in natural language processing contexts. Given a set of documents, LDA uses word count distributions to define a set of topics that each document discusses. For example, documents may primarily discuss politics, cooking, or sports. Gibbs sampling is used to learn unlabeled topics by a weighted vector of vocabulary words. Simultaneously, documents are assigned weighted vectors of topic distributions. The resulting topic vectors as defined by vocabulary are unlabeled, however, by interpreting those words with high weight in a given topic, one may assign a label to the topic heuristically. For example, one topic may contain high weights for president, prime minister, counselor, and court room, while another may contain high weight for the words spatula, bruschetta, and baguette. A human interpreter could see these patterns and call the first topic “politics” and the second topic “cooking”.

The Dirichlet distribution is commonly used to model microbiome data ***Holmes et al. (2012); Harrison et al. (2020); Wadsworth et al. (2017)***, and LDA in particular has proven to be a powerful and robust tool for modeling 16S data in various environments ***Knights et al. (2014); Sankaran and Holmes (2018); Deek and Li (2021); Breuninger et al. (2021); Sommeria-Klein et al. (2020); Okui (2020)***. LDA is a parametric form of dimensionality reduction; it has the advantage of additional power as compared to other non-parametric forms like PCoA, GloVe, or transformer neural networks ***Chin and Lee (2008)***. In the case of multi-omic analysis, where data representing multiple omic measurements on the same samples is very limited, parametric methods are particularly useful. LDA allows for mixed membership, so a sample can be represented by multiple subcommunities of bacteria, genes, or metabolites, making it more appropriate for capturing complex phenomena.

### Value of cross-omic microbiome topics

Cross-omic topics represent gut microbiome landscapes that can be viewed through the lens of individual omics, but that are present regardless of the lens. We are not the first to consider microbial landscapes as a basis for comparing samples.

Perhaps the most commonly cited example of considering microbial landscapes is that of en-terotypes. In a 2011 paper, Arumugam et. al introduced the idea that the human gut can come in three varieties, those dominated by Bacteroides, Prevotella, and Ruminococcus genera respectively. Laterwork by Knights et. al suggested that the enterotype distribution is actually continuous rather than discrete ***Knights et al. (2014)***. Most recently, Symul et. al consider how we might use approaches like topic modeling to identify sub-communities of samples within this continuous microbiome space in the vaginal ecosystem ***Symul et al. (2022)***. To our knowledge, the approach of topic modeling to define microbial sub-communities has been reported only for 16S data, with other omic data as supporting evidence. Here, we illustrate that the same sub-community or subprocess structures can be observed through the lens of multiple levels of biology (e.g. community membership, genomic function, and metabolic result), and that the sub-processes (topics) identified by each omic do correspond significantly with one another to define sub-processes across omics.

By comparing only samples with similar community structure, and only then identifyingthe spe-cificfactors by which they differ, our results become more meaningful. For example, in macroecology, it is standard practice to compare population sizes between species in similar environments only, as comparisons between vastly different habitats may be correct, but not meaningful. In microbial ecology, we often perform differential abundance analysis to identify features that differ between host phenotypes without consideration for the broader microbial landscape. We argue that it is not enough that samples come from the same tissue (e.g. gut), we must also consider the ecology between individuals as a relevant factor. One approach to accomplish this grouping of similar samples is to cluster samples based on similarities of diet and/or host genotype, however, the effect these factors have on the microbial environment is not fully understood, and does not account for all differences observed in broad microbial landscapes. Comparisons made on the basis of dietary similarities are also usually subjective to self-report bias. Instead, methodologies that act on the microbial landscapes themselves promise to be more consistent across studies and inherently mitigate bias derived from an incomplete understanding of microbiome-diet relationships.

Additionally, topics may represent underlying metabolic processes that are independently informative of microbial ecology principles. In the “Interpreting Topics by Feature Weights” section, we observe that biologically related features (ex. sets of dinucleotides or genes utilized by bacteria in stressful environments) are all attributed to the same topic. This consistency indicates that biological relationships may be captured in this methodology, and suggests that topic modeling may be used to broaden our understanding about gene, metabolite, and taxonomic species relationships. We found that topics learned in each omic dataset independently significantly correlated with other topics from other omics, which was not true of null simulated data. We hypothesize that this implies some universality of the processes represented by cross-omic topics, however, further research is needed to validate these topic distributions on independent datasets.

One benefit of identifying sub-processes based on multiple omics is mitigation of bias. Each omic technique provides useful information about one level of biological processing, but each is inherently incomplete and subject to unique sources of bias during processing. 16S data acts as a census of the bacterial species represented in the community, and their relative abundances. Because it involves an amplification step, it represents a deeper sampling of the microbial diversity present, but is also subject to amplification bias ***McLaren et al. (2019a)***. It also has very limited capacity to represent function or metabolism within the bacterial community ***McLaren et al. (2019b)***. Shotgun metagenomic (MTG) data provides a shotgun representation of the community gene pool.

Because it lacks an amplification step, it represents only the most common genes. It also represents only genetic potential, as opposed to those genes undergoing expression or translation ***Quince et al. (2017)***. Shotgun metatranscriptomic (MTT) data provides a shotgun representation of the community translation pool. Like MTG, it represents only the most common genes, and can be additionally highly variable between timepoints ***Shakya et al. (2019)***. Metabolomic (MBX) data provides a measurement of the metabolites and relative concentrations present in an environment. Metabolites may originate from bacteria or host, and data is additionally limited to those most common metabolites ***Johnson and Gonzalez (2012)***. Individually, all are subject to unavoidable bias, however, integrative multi-omic approaches such as topic modeling have the potential to mitigate these biases - phenomena that are represented in all data modalities are more likely to be ubiquitous, universal, and consistent phenomena.

### Dietary habits as related to cross-omic microbiome topics

It is commonly accepted that diet influences microbiome composition and function and vice versa, and it is not known explicitly what dietary habits promote the healthiest microbiomes. We found that correlations between dietary variables and cross-omic topics, which represent the relationship strength between specific dietary practices and microbiome composition and function, spontaneously group dietary choices into two groups. Group1 consists of fruit, vegetable, fermented vegetables, meat, seafood, home prepared meals, and vitamins, while group2 consists of sugary and starchy foods, sweetened drinks, dairy, and ready to eat meals. Interestingly, self-assembled groups clearly fall in line with current pre-determined dietary health standards ***Breuninger et al. (2021)***.

In this case, it is impossible to distinguish between effects on the microbiome driven by lack of an important “healthy” food group vs. presence of an “unhealthy” food group. Further studies may be designed to include participants actively eating all the necessary food groups in addition to some unhealthy options, as well as participants lacking some healthy food group choices to understand whether presence or absence of healthy eating choices is more influential on the gut microbiome.

### Cross-omic microbiome topics as defined by microbial functions

#### Topic1: healthy/general phenomena

We hypothesize that Topic1 may represent base metabolism functionalities across many forms of life, including beta-oxidation (metabolism of fats), cytochrome functionality, methane metabolism, and ion transport. This is supported by a strong correlation between Topic1 values and 16s-based alpha diversity metrics, which are strongly linked to increased resilience and general healthy status ***Xu and Knight (2015); Manor et al. (2020); Hagerty et al. (2020); Menni et al. (2017); Fassarella et al. (2021)***. Topic1 correlates most strongly with dietary variables such as fruit, vegetable, fat/oil, home prepared meal, meat, seafood, and vitamin consumption frequency, which are largely in line with the Healthy Eating Index “adequacy components” (eating more corresponds to a healthier diet) ***Arem et al. (2013)***.

Within the MBX topics, there are 3 that belong to cross-omic Topic1. The first is represented by high values of 18 carbon, 3 times unsaturated lipids such as anacardic acid, heptadecatrienoate, and 18:3 glycerols. The second is represented by higher values of metabolites related to CYP1A2 metabolism of caffeine such as 1,3,7 trimethylurate, AAMU (CYP1A2 caffeine metabolism products), and alpha tocopherol acetate (CYP1A2 inhibitor) ***Labedzki et al. (2002); Nyéki et al. (2003); Le Marchand et al. (1997)***. We hypothesize that this topic may represent the products of CYP1A2 activity of the host, which may be modified with caffeine consumption (in the form of coffee, tea, or chocolate), as well as by cruciferous and apiaceous vegetable consumption ***Lampe et al. (2000); Tantcheva-Poór et al. (1999)***. The last is represented by high values of PAHSA and ximenoylcarnitine, long chain and very long chain fatty acids that relate to mitochondrial function. Specifically, PAHSAs are known for their anti-inflammatory and insulin controlling action, while ximenoylcarnitine is less studied, but as an acylcarnitine likely has to do with beta-oxidation activity in the mitochondria ***Brejchova et al. (2020); Schultz Moreira et al. (2020)***. There are two MTG topics that cluster into cross-omic Topic1. They are characterized by ubiquitous functions like glutamine and serine/threonine kinases and by environmental adaptation genes like heavy metal and drug exporter pumps ***Blanco et al. (2016); Pereira et al. (2011)***. There are two MTT topics that cluster into cross-omic Topic1 as well. They are characterized by high values of methyl-coenzyme M subunits used in methane metabolisms, as well as eukaryote-specific genes such as yeast plasma membrane ATPase, and yeast amino acid transporter. This same topic is also high in ubiquitous functions like eukaryotic translation initiation factors, pyruvate decarboxylase, heat shock protein, and MFS sugar transporters ***Henderson and Maiden (1990)***. While the methane metabolism transcripts most likely represent archaea activity in the gut ***Gaci et al. (2014)***, the second MTT Topic1 more likely relates to eukaryotic metabolisms, specifically yeast. Contributing to this conclusion are the eukaryotic specific transcription factor and yeast specific genes that define this topic, as well as the fact that all transcripts mapping to the human genome were removed (see Methods), eliminating the main source of eukaryotic gene transcripts. Lastly, we see some Ruminococcaceae species along with unidentified Clostridia members highly represented, that, when combined with the high vegetable associations and hypothesized methane metabolisms, imply prevalent fermentation processes. ***Ze et al. (2012)***.

#### Topic2: age-associated phenomena

In contrast, Topic2, which is also significantly inversely correlated with age, correlates with low quantities of the above food groups, and high quantities of starchy, sugary, pre-packaged and restaurant foods, which are largely in line with the Healthy Eating Index “moderation” components (eating less corresponds to a healthier diet) ***Arem et al. (2013)***. In MBX Topic2, we see high values of steroid sulfates, specifically androstenediol mono and di sulfate, tetrahydrocortisol sulfate, and pregnenediol sulfate, DHEA sulfate, and alpha -CEHC sulfate. These steroid compounds are responsible for physiological changes seen during puberty, but are inactive in their sulfonated forms ***Mueller et al. (2015)***. Thus, we hypothesize that Topic2 may capture the effect of age with an inverse relationship. While steroid sulfates are not considered actively hormonally, some do act as neurosteroids, which can modulate GABA and NMDA receptors among other targets ***Gibbs et al. (2006)***. Steroid sulfatase is a potential modifier of cognition in attention deficit hyperactivity disorder ***Stergiakouli et al. (2011)***. It is suggested that STS dysfunction (too many sulfates on hormones) predisposes an inattentive subtype of ADHD ***Brookes et al. (2008)***. We did not find Topic2 to be associated with Autism severity, and it was not able to distinguish between autistic and TD individuals (data not shown).

In Topic2 MTG, we observed high values of a key enzyme in steroid hormone synthesis, phos-phomevalonate kinase, which is consistent with the steroid metabolites observed in the MBX data ***Cerqueira et al. (2016)***. We also observe high values for genes used in bacterial horizontal gene transfer, namely competence proteins and transposases. Upregulation of these gene sets has been observed from bacterial in stressful environments, and diets heavy in starch and sugars can affect biofilm formation as well as competence ***Solomon and Grossman (1996); Klein et al. (2009); Ogura et al. (1999) Klein et al. (2009); Ogura et al. (1999); Cordero et al. (2022)***. Topic2 also represents high values Ruminiclostridium species, Blautia and Tyzzerella and Faecalitalea genera members, many of which are associated with chronic inflammation. Ruminiclostridium is considered a potential pathogen associated with obesity and inflammation, and has been found to be increased in younger individuals in macaques, dairy cows, and humans ***Zhang et al. (2020); Duan et al. (2019); Zhang et al. (2019)***. Tyzzerella was found to be profoundly overrepresented in Crohn’s disease patients, increased in patients with high cardiovascular disease risk profiles, and correlated with lower healthy eating scores ***Olaisen et al. (2021); Kelly et al. (2016); Liu et al. (2019)***. It should be noted that while Topic2 was strongly inversely related to age, participants were no younger than 2 years old and were not being breast-fed, and so Topic2 is not representative of a newborn gut microbiome. The precise relationship between the observed competence proteins, pro-inflammatory genera, and age remains to be elucidated.

#### Topic3: transcriptional regulation phenomena

In Topic3, we see high values of dinucleoside monophosphates and genes for RNA polymerase subunits, elongation factor G, and HSP20. Dinucleoside monophosphates have been recently suggested to function as RNA caps in bacteria, to either initiate transcription or protect against RNA degradation ***Olaisen et al. (2021); Hudeček et al. (2020)***. Alternatively, they may also be an artifact of metabolomics pipeline processing. In 16s data, we see high values for two species, one from the family Christensenellaceae and the other from Odoribacter. MTG and MTT data do not present strong feature attributions for Topic3, however, we do see that the overall topic weight across samples attributed to Topic3 is incredibly high (Supp. Figure 6), supporting the interpretation of Topic3 as transcription related factors.

#### Topic4: opportunistic pathogenesis phenomena

Topic4 is characterized less strongly by metadata variables, but still represents lower intake of vegetable, fruit, fat/oil, meat, and home prepared meals. In MBX data, we see high values of sulfates, specifically catechol, dihydrocaffeate, and furaneol sulfates. Catechol is used as a pesticide and as a precursor to many chemical products, can be manufactured by multiple bacterial species including those found in the human gut ***Balderas-Hernández et al. (2014)***, and has been found to have both anti-bacterial and anti-fungal action ***Kocaçalişkan et al. (2006)***. Dihydrocaffeic acid is produced during colonic fermentation of wheat ***Koistinen et al. (2022)***. Interestingly, although children ages 2-7 are unlikely to be consuming coffee, all three compounds are known to increase upon coffee consumption ***Goldstein et al. (2021); Tressl et al. (1978)***. Additionally, metabolites that reflect beta-oxidation processes in the host are increased in MBX data, specifically, myristoylcarnitine, linoleoylcarnitine, 7-ketolithocholate, 3-hydroxyisobutyrate, and 2-O-methylascorbic acid. Acylcarnitines along with 3-hydroxyisobutyrate are intermediates of beta oxidation ***Rutkowsky et al. (2014); Nilsen et al. (2020)***, while 7-ketolithocholate and other bile acids contribute to fat absorption and may influence the availability of fatty acids for catabolism ***Ahmad and Haeusler (2019)***. 2-O-methylascorbic acid is understudied in the literature, however, 2-O-ethyascorbic acid, or Vitamin C, is a cofactor for carnitine biosynthesis which is necessary for beta-oxidation ***Rebouche (1991)***. Interestingly, bacteria may also use carnitine as an osmoprotectant to increase thermotolerance, cryotolerance and barotolerance, and may use 3-HB as a substrate for the synthesis of polyhydroxybutyrate, which is a reserve material ***Meadows and Wargo (2015); Nilsen et al. (2020)***. High concentrations of these metabolites contribute to the conclusion that Topic4 may represent a stressful or challenging environment for many microorganisms, and may increase the opportunity for opportunistic pathogen growth. In MTG, we see a strong contribution from genes associated with pathogenesis and metabolism in stressful environments. Autotransporter family proteins are often associated with virulence functions such as adhesion, aggregation, invasion, biofilm formation and toxicity, usher proteins facilitate pillus formation, and adhesin/invasin can induce bacterial aggregation and biofilm formation ***Wells et al. (2007); Remaut and Ben-Tal (2014); Sherlock et al. (2005)***. Miniconductance mechanosensitive channel confers protection against mild hypoosmotic shock ***Schumann et al. (2010)*** and bacteriophage proteins signal a stressful environment for commensals. In MTT, we see similarly high values of universal stress protein E, outer membrane usher protein, biofilm regulator BssR, as well as regulatory proteins RseB, molecular chaperone IbpA, and chromosome partition protein MukB. Many of these genes are upregulated during microbial adaptation to stressful environments, and by opportunistic pathogens in particular ***Siegele (2005); Domka et al. (2006); Miwa et al. (2021)***. Lastly, 16s data shows particularly high values for a Veillonella species, as well as for species from the Blautia, Faecalibacterium, Intestinibacter, and Anaerostipes genera. Veillonella species are the strongest contributors to Topic4; they are well known for their behavior as opportunistic pathogens in the human gut and dental plaque and exhibit strong adhesion and biofilm formation capacities mediated by autotransporters ***Béchon et al. (2020)***.

### Subsetting population by topics reveals metabolic differentials between ASD and TD cohorts-neurotransmitters and fatty acids

In cluster1 (high topics 1 and 3), which corresponds to largely healthy eating habits including high quantities of vegetables, fruits, and home prepared meals, we found autistic individuals to have lower stool abundances of neurotransmitter precursors carboxyethyl GABA, glutamine, and tryptophan. Autism is characterized by complex neurobehavioral and neurodevelopmental criteria including social interaction, restricted and repetitive behavior, and altered sensory processing ***Lord et al. (1989***, 2000). Many have reported dysregulation in the glutamate-glutamine cycle in both plasma and brain tissues resulting in altered concentrations of glutamine and GABA in people with Autism ***Al-Otaish et al. (2018), Marotta et al. (2020), Cochran et al. (2015), Horder et al. (2013)***. Likewise, multiple components of the tryptophan metabolism pathway are neuroactive, including serotonin, kynurenine, and quinolinic acid ***Oxenkrug (2013); Lapin (1998)***. Dysregulation in tryptophan metabolism is hypothesized as a major contributing agent to the gut-brain axis, and is associated with Autism ***Adams et al. (2011)***. Tryptophan may also become NAD+ through the kynurenine pathway, where dysregulation may imply issues with metabolic regulation ***Castro-Portuguez and Sutphin (2020)***. While CEGABA is less studied, evidence suggests that it is active in the central nervous system and may strengthen cortical inhibition and actdirectlyon the lower brain stem ***Adams et al. (2011); Bo et al. (1988)***. It also demonstrated anti-convulsant activity in guinea pigs ***Savoldi et al. (1987)***.

In cluster2 (high topics 2 and 4), which corresponds to largely unhealthy eating habits including high quantities of sugary and processed food and drinks and low quantities of fruits and vegetables, we found autistic individuals to have lower stool abundances of fatty acids, acylcarnitines, quinolinediol, and 3-hydroxy-5-cholenoic acid. Mitochondrial dysfunction, as well as fatty acid and acylcarnitine concentrations in serum, has been implicated in subgroups of autistic individuals ***Barone et al. (2018); Jay Gargus and Imtiaz (2008); Rossignol and Frye (2012)***. Concentrations of bile acids like 3-hydroxy-5-cholenoic acid have also been implicated in Autism, perhaps in part because bile acid concentrations directly influence the absorption and therefore availability of fatty acids ***Golubeva et al. (2017); Wu (2017)***. We were unable to find reference to quinolinediol in ASD literature, however, quinolinic acid, a related compound, was found to be decreased in cerebrospinal fluid of 12 children with autism ***Zimmerman et al. (2005)***. Quinolinic acid also serves as a precursor to NAD+, and this pathway may be used as an alternative to synthesize NAD+ in the context of oxidative stress as would be seen during mitochondrial dysfunction ***Sahm et al. (2013)***. Mitochondrial dysfunction and oxidative stress has been reported in only 5% of autistic individuals, however, this is much higher than the expected 0.01% in the rest of the population ***Sahm et al. (2013); Rossignol and Frye (2012)***.

Both neurotransmitter imbalances and metabolic dysregulation have been implicated in Autism literature, depending on the study. This study implicates both factors depending on the subtype of the gut microbiome, which itself relates to both diet, microbial gene content and expression, and metabolic environment. This successful subtyping, along with correlations between some topics and specific behavioral metrics, lead us to suggest that the gut microbiome may be effective at identifying subgroups of Autism, and may give some indication as to the metabolic differences seen between those subgroups of autistic and typically developing children.

## Conclusion

Topic analysis revealed generalized microbial processes consistent across multiple profiling methods. Topics correlate with specific dietary variables, and we hypothesize that each represents a combination of microbial phenomena, specifically healthy/general function, age-associated function, transcriptional regulation, and opportunistic pathogenesis. Furthermore, topic distribution can be used to cluster samples such that samples in the same group share similarities in their multi-omic representation, which subsequently implicates both neurotransmitter precursors and fatty acid derivatives in Autism. Future multi-omic datasets along with *in vitro* studies promise to provide further insight into the consistency and accuracy of topic definitions. Confirmation of these topics could lead to new and improved avenues from human gut microbiome engineering and microbiome-targeted therapeutics.

## Materials and Methods

### Cohort and Metadata

Stool samples from 196 children between ages 2-7 from all over the United States were collected via a crowdsourced initiative. Children were all from families with two siblings, one previously diagnosed with Autism by a health care provider and one typically developing. Dietary, lifestyle, demographic, and host health information were reported by parents of guardians via a questionnaire for each child (see ***Tataru et al. (2021)***). Parents and guardians also filled out a Mobile Autism Risk Assessment (MARA) to document autism-associated behaviors in their children with the Autism diagnosis ***Duda et al. (2017)***. Samples were then subjected to multi-omic sequencing. In total, 81 of the stool samples received all four omic measurements, and these data were used in defining cross-omic topics. However, some individuals provided multiple stool samples over time, and in topic modeling on each omic independently, all available data was used: 16S (456 stool samples across 152 individuals), MTG (193 stool samples across 186 individuals), MTT (178 stool samples across 178 inviduals), and MBX (175 stool samples across 175 individuals.

### Stool Collection and Storage Stool

Samples were collected by the parents using a preservative buffer (Norgen Biotek, ON, Canada). We collected two samples per child: one sample was preserved at room temperature in a preservative buffer, and the second one was collected in a tube without a preservative buffer but immediately frozen at home at −20. This frozen sample was shipped back overnight with two ice packs provided to the participants, while the samples in preservative were shipped at room temperature. Once received, stool samples were stored at-80C until processing.

### Profiling Techniques

#### 16S Sequencing

DNA was extracted using the Qiagen MagAttract PowerMicrobiome DNA/RNA Kit according to manufacturer’s guidelines. DNA was quantified using the Qubit^®^ Quant-iT dsDNA High Sensitivity Kit (Invitrogen, Life Technologies, Grand Island, NY). The V4 regions of the 16S rRNAgene were amplified using primers as described in ***Caporaso et al. (2012)***, and PCR products were quantified by fluorometric method (Qubit or PicoGreen from Invitrogen, Life Technologies, Grand Island, NY). E uimolar amounts of amplicons were mixed and sequenced using Illumina MiSeq for 250 cycles at Second Genome. All samples from the same families were sequenced in the same batch. Samples with fewer than 20,000 reads were re-sequenced up to three times.

#### Shotgun Metagenomic (MTG) and Metatranscriptomic (MTT) Sequencing

The MTG library was constructed with the same DNA extracts as 16S-V4. For MTT, DNA extraction and digestion was performed using Qiagen MagAttract PowerMicrobiome RNA Kit and Invitrogen TURBO DNA free kit according to manufacturer’s guidelines. rRNA depletion was also performed using Ribo-Zero Gold rRNA Removal Kit (Epidemiology). All samples were prepared for sequencing with the Illumina NexteraXT kit and quantified with Quant-iT dsDNA High Sensitivity assays. Libraries were pooled and run with 150 bp paired-end sequencing protocols on the Illumina NextSeq 550 platform.

#### Metabolomics

Untargeted metabolomics was performed on the preservative-free stool samples by Metabolon, Inc. Samples were extracted with methanol to precipitate protein and dissociate small molecules bound to protein or trapped in the precipitated protein matrix, followed by centrifugation to recover chemically diverse metabolites. The resulting extract was divided into five fractions: two for analysis by two separate reverse phase (RP)/UPLC-MS/MS methods using positive ion mode electrospray ionization (ESI), one for analysis by RP/UPLC-MS/MS using negative ion mode ESI, one for analysis by HILIC/UPLC-MS/MS using negative ion mode ESI, and one reserved for backup. Ultrahigh Performance Liquid Chromatography-Tandem Mass Spectroscopy (UPLC-MS/MS) was performed. All methods utilized a Waters ACQUITY ultra-performance liquid chromatography (UPLC) and a Thermo Scientific Q-Exactive high resolution/accurate mass spectrometer interfaced with a heated electrospray ionization (HESI-II) source and Orbitrap mass analyzer operated at 35,000 mass resolution. The sample extract was dried then reconstituted in solvents compatible with each of the four methods. Each reconstitution solvent contained a series of standards at fixed concentrations to ensure injection and chromatographic consistency. One aliquot was analyzed using acidic positive ion conditions, chromatographically optimized for more hydrophilic compounds. In this method, the extract was gradient-eluted from a C18 column (Waters UPLC BEH C18-2.1×100 mm, 1.7 μm) using water and methanol, containing 0.05% perfluoropentanoic acid (PFPA) and 0.1% formic acid (FA). A second aliquot was also analyzed using acidic positive ion conditions, but was chromatographically optimized for more hydrophobic compounds. In this method, the extract ias gradient eluted from the aforementioned C18 column using methanol, acetonitrile, water, 0.05% PFPA and 0.01% FA, and was operated at an overall higher organic content. A third aliquot was analyzed using basic negative ion optimized conditions using a separate dedicated C18 column. The basic extracts were gradient-eluted from the column using methanol and water, however with 6.5mM Ammonium Bicarbonate at pH 8. The fourth aliquot was analyzed via negative ionization following elution from a HILIC column (Waters UPLC BEH Amide 2.1×150 mm, 1.7 μm) using a gradient consisting of water and acetonitrile with 10mM Ammonium Formate, pH 10.8. The MS analysis alternated between MS and data-dependent MSn scans using dynamic exclusion. The scan range varied slightly between methods, but covered approximately 70-1000 m/z.

### Data Processing

#### 16S

DADA2 (Callahan et al. 2016) was used to generate Amplicon Sequence Variants (ASVs). Raw sequence reads were processed applying default settings for filtering, learning errors, dereplication, ASV inference, and chimera removal. Truncation quality was set to 2. Ten nucleotides were then trimmed from each terminus of each read, both forward and reverse. ASVs were mapped to an in-house strain database: StrainSelect (StrainSelect, http://strainselect.secondgenome.com/, version 2019) using USEARCH (Edgar 2018). StrainSelect is a repository of monikers for known isolated microbial strains, their various synonyms and their genomic sequence identifiers. A DNA sequence observed from clinical material was assigned a strain-level annotation only when it uniquely matched one and only one strain. 6,150 ASVs were generated from this process.

#### MTG and MTT

Reads were trimmed for adapter sequences and low-quality ends with Trimmomatic (Bolger, Lohse, and Usadel 2014), then contaminant sequences were removed with Bowtie2 (Langmead and Salzberg 2012). Host sequences were identified and removed with Kraken (Wood and Salzberg 2014). For MTT data, an additional step for rRNA removal was performed using SortMeRNA 2.0 (Kopylova, Noé, and Touzet 2012) prior to the host sequence removal. Filtered DNA sequences from MTG and MTT were mapped against a reference database of proteins within the KEGG (May 2019 release) and hits that spanned more than 20 amino acids with more than 80% similarity were collected. A total of 10,543 KOs were observed in MTG data, and 10,625 KOs were detected in MTT data.

#### MBX

Raw data were extracted, peak-identified, and quality control (QC) processed using Metabolon’s hardware and software. Compounds were identified by comparison to library entries of purified standards or recurrent unknown entities. Peaks were quantified as area-under-the-curve detector ion counts. For studies spanning multiple days, a data adjustment step was performed to correct block variation resulting from instrument inter-day tuning differences, while preserving intraday variance. 1,267 metabolites were identified, and 1,025 annotated metabolites were used in the downstream analysis.

### Data Analysis

#### Normalization

To select a normalization method, we sought to minimize the within sibling to between sibling distance ratio using manhattan, euclidean, canberra, clark, bray, kulczynski, jaccard, and altGower distances from the vegan package. We found that DeSeq2 normalization minimized the ratio for all of the distance metrics in 16S data, and the highly related Relative Log Expression (RLE) normalization minimized the ratio for MBX data especially. Differences in the sibling ratio for MTG and MTT were slight, and so RLE was selected for these data as well for the sake of consistency (Supp. Figure 9). The R packages DeSeq2 and edgeR were used to normalize data respectively.

The high counts of the MTG, MTT, and MBX datasets made the Gibbs samples strategy for topic model training too computationally expensive. Therefore, we log transformed the counts of each dataset after normalization and before model training.

### Topic modeling

We performed Latent Dirichlet Allocation (LDA) as a form of dimensionality reduction and multiomic integration. LDA is a generative statistical model traditionally used in natural language processing that models documents as deriving from a set of topics, and topics as deriving from a set of words. In our case, we treat samples as documents and microbial features as words. As such, each sample is defined by a probability vector over K possible topics, and each topic is defined by a probability vector over V possible microbial features (e.g. ASVs, KOs, metabolites).

Under LDA, to generate each word in a document (*w_d_n__*), first get that document’s (d) topic probability vector (*θ_d_*) and select a topic (*z_d_n__*) by drawing from a multinomial. Then, get that topic’s word probability vector (*β_z_d_n___*), and select a word (*w_d_n__*) using a multinomial. Each of the topic probability vectors is modeled as a Dirichlet distribution over hyperparameter *α*, and each of the word probability vectors is modeled as a Dirichlet distribution over a different hyperparameter *γ*. After model fitting, the Θ and *β* matrices can be accessed to obtain the topic distribution per document and the word distribution per topic. In this case, we access these matrices to obtain the topic distribution per sample and the microbial feature distribution per topic (Figure 1D). Models were trained using the topicmodels package in R.

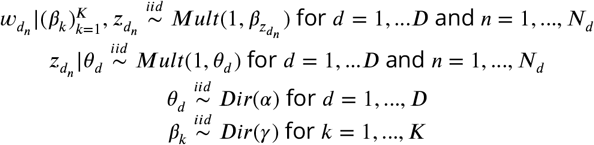

### Selecting number of topics per model

Models for each omic were trained independently, and the number of topics was selected per model. To evaluate model fit, we simulated sample count data drawn from each model, and compared it to the true distributions observed. We used three metrics, correlation of quantiles of sample distributions, correlation of quantiles of feature distributions, and correlation of pairwise marginal distributions (correlation of feature correlations). For each omic, we selected the minimum number of topics that produced the highest correlation across all of the three metrics. The final number of topics selected were 4 for 16s data, 5 for MTG data, 4 for MTT data, and 7 for MBX data (Supp. Figure 10).

### Identifying cross-omic topics

To determine the optimal number of cross-omic topics, we selected the minimum number of topic clusters that minimized the gap statistic amongst clustered topics. First, we clustered topics into n groups based on their spearman correlation across samples. Then, for each cluster, we calculated the average distance between topics in that cluster (within group distance), and divided by the average distance between topics in that cluster and all other topics (between group distance). We used manhattan, euclidean, and inverse correlation as distance metrics, and selected the minimum number ofclusters that minimized the above ratio adequately. In the end, we selected 4 cross-omic topic clusters (Supp. Figure 1 A). Topics were visualized by PCoA on the topic distribution across samples using manhattan distance (Supp. Figure 1 B).

We additionally found that single-omic topics within a cross-omic topic group significantly cor-related with one another far more frequently than would be expected by random chance. To test this, we created 15 null count tables, drawing each sample from a multinomial where probabilities are assigned for each feature using a random sample’s actual feature distribution. We then fit an LDA model to each of the null count tables using the same process, parameters, and random seeds as were used in the original models. We then clustered the topics produced from the null models into four clusters, as was done with the original data, and counted the number of significant (spearman correlation test, p < .05) correlations existed between topics that shared the same cluster. We found that null topics in the same cluster shared very few significant correlations and null topic distributions across 15 permutations had correlation structures stronger than the true data (Supp. Figure 1 C). We additionally visualized a correlation matrix between null topics (Supp. Figure 1 D) and between true topics (Supp. Figure 1 E).

It is relevant to note that the treatment of MTT topics differed slightly from the rest. The total attributable weight across all samples to topic3 MTT was orders of magnitude higher than the other MTT topics (Supp. Figure 6). In addition, no specific features differentiated topic3 MTT from the other MTT topics (data not shown). Because this over attribution of topic3 MTT was not informative of differences between samples nor interpretable, and because its inclusion swayed all downstream clustering analyses considerably, we removed this component.

### Interpreting topics by feature weights

We used the feature weight vectors (*β*)s to interpret each cross-omic topic. We limited features to those with high standard deviations across topics, as we wanted to identify features that were specific to certain topics and not ubiquitous across topics. Specifically, we identified features where the max value was less than 1.5 standard deviations over the median. We also limited features to those with high values along axes of maximal variation between topics, as these are the features driving topic definitions and separation most strongly. Specifically, we performed singular value decomposition on the topic by feature weight matrices independently (including all features), and recorded features with weights along the PCA axes above the 99th percentile for 16s and MBX data, and above the 99.9th percentile for MTG and MTT features. Features that fulfilled both of the above criteria are reported in Figure 2. These thresholds were selected heuristically based on the maximum number of features that could be clearly visually represented.

### Clustering samples by topic distribution

To cluster samples by their topic distribution, we used the R package pheatmap, which performs hierarchical clustering using inverse correlation as a distance metric.

### Differential analysis

Differential analysis was performed to determine the association between any given microbial feature between the Autism and typically developing cohorts after clustering. First, we selected likely candidates using the Boruta package in R. The method performs a top-down search for relevant features by comparing original attributes’ importance with importance achievable at random, estimated using their permuted copies, and progressively eliminating irrelevant features to stabilize that test ***Kursa and Rudnicki (2010)***.

In more detail, filtering features with Boruta works as such: 1) create an extended dataset by adding columns of randomly permuted features; 2) train a random forest classifier on the extended dataset to predict phenotype (Autism vs. typically developing) and calculate feature importance as the mean decrease in accuracy; 3) if the true feature Z score is higher than the maximum Z score of it’s permuted versions over 100 runs, consider this feature “important”. To increase consistency and generalizability, we repeated this described test 100 times, and tallied the number of times any given feature was considered “important”. Features in the top 90th percentile of tallied counts were passed to a subsequent Wilcoxon rank sum, and were reported if their differential was significant (p<.05) after Benjamini & Hocherg correction ***Benjamini and Hochberg (1995)***.

Number of input features for each omic and number of features that passed the first step of Boruta filtering are reported here: 1187 features input and 7 features passed filtering (MBX), 10543 features input and 15 features passed filtering (MTG), 10625 features input and 8 features passed filtering (MTT), 5265 features input and 4 passed filtering (16S).

## Data Availability

All analyses and processed data files are available at: https://github.com/MaudeDavidLab/multiomics_topic_modeling. Data are also available at https://files.cqls.oregonstate.edu/David_Lab/M3/. Raw data available upon request. This study was authorized by the Institutional Review Board protocol number 30205.

---

**Supp. Figure 1.**
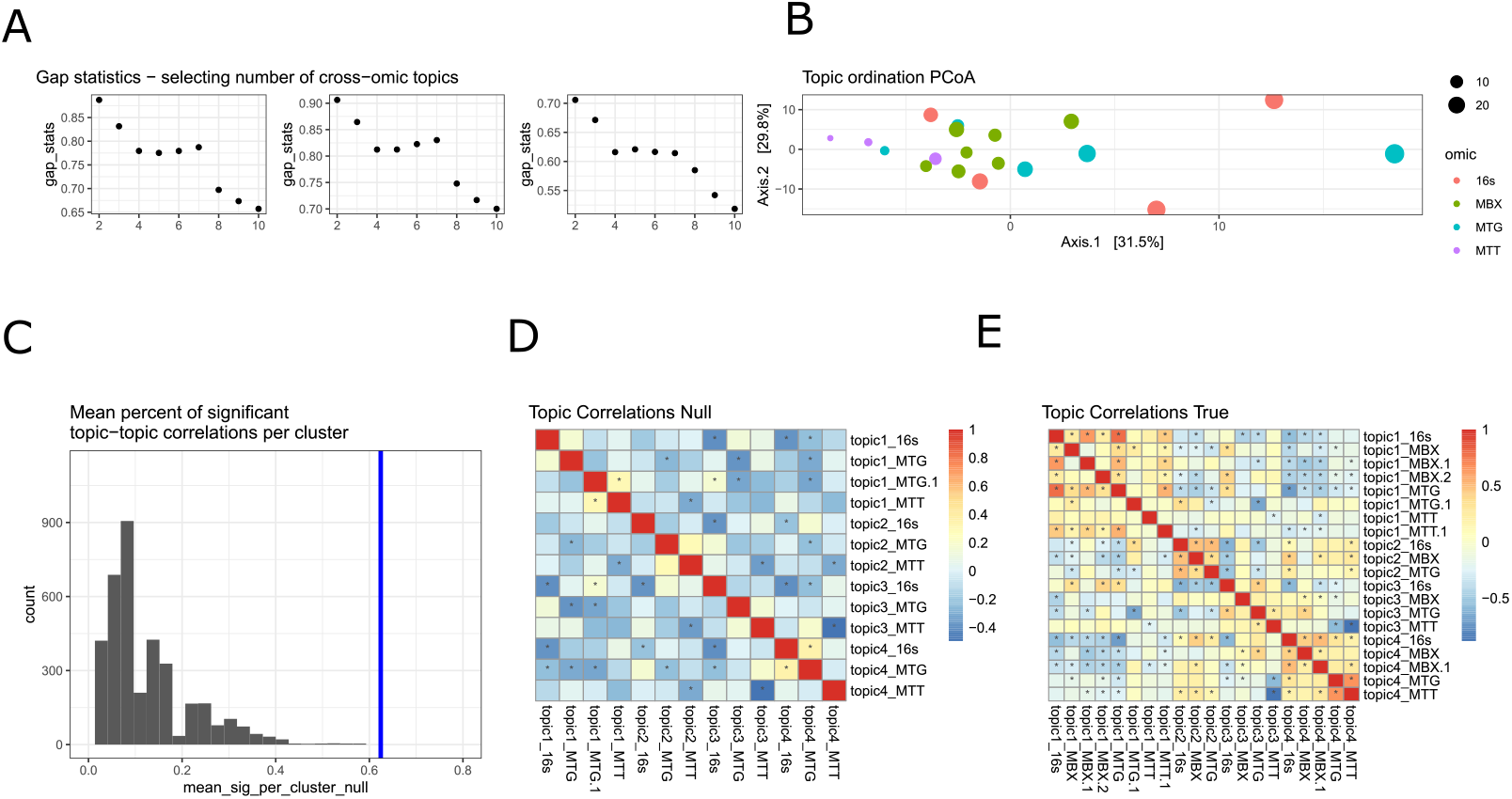
Cross-omie topic evaluation. Gap statistics (within/between cluster distances) were calculated using manhattan, euclidean, and inverse correlation distance metrics for 2-10 cross-omic topics. The elbow method was used to select 4 cross-omic topics (A). Topics were visualized by PCoA on the topic distribution across samples using manhattan distance. Size of points represents the total weight attributed to a topic across all samples (how common the topic is) and color represents omic of origin (B). Topic-topic correlations within a cross-omic topic cluster are significantly more correlated than would be expected by random chance. The mean number of significant (spearman correlation p<0.05) topic-topic correlations within a cluster using null simulated data (15 permutations) is between 0 and 0.4. The true number mean of significant correlations is over 0.6 (C). Topic-Topic correlation matrix using null simulated data (D). Topic-Topic correlation matrix using true data (E). Cells are colored by spearman correlation coefficient and star indicates significant (p<0.05). Multiple topics from the same omic dataset may be in the same cross-omic topic, denoted by”.1” or”.2”, etc.

**Supp. Figure 2.**
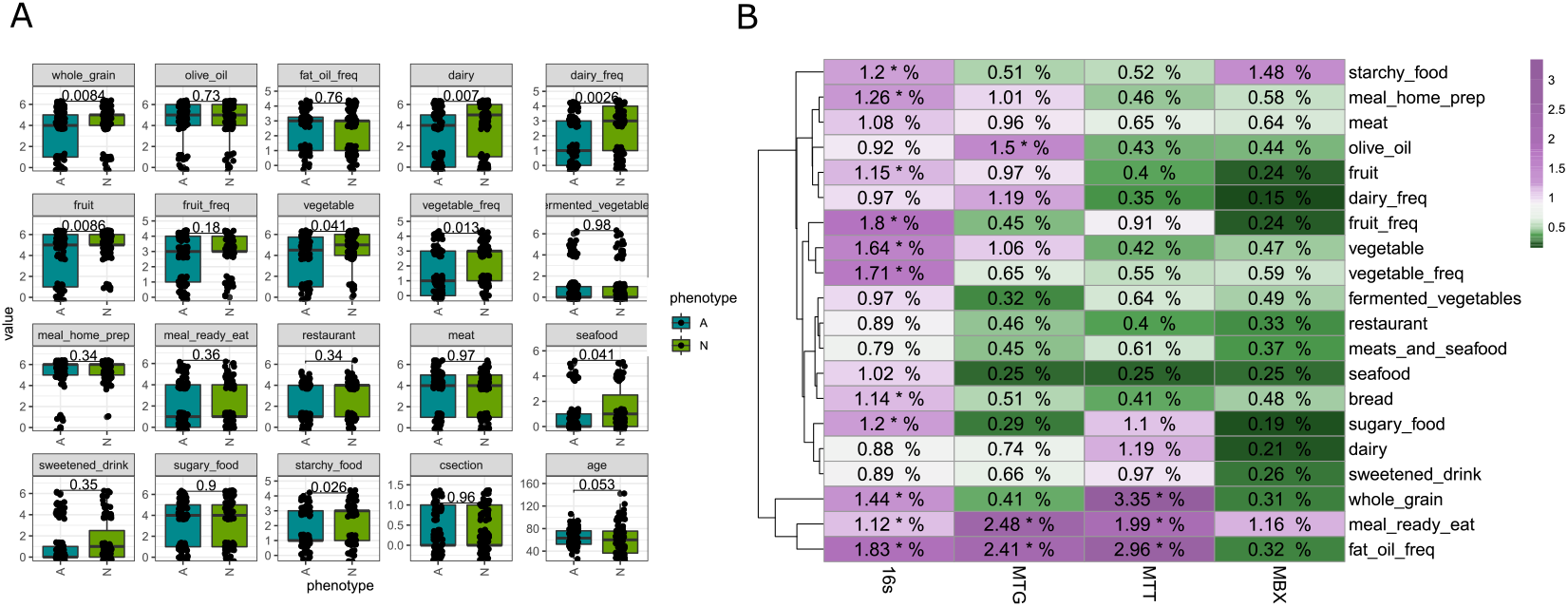
Dietary and lifestyle characteristics between autistic and typically developing children compared using a Wilcox rank sum test (A). Association between individual dietary variables and omic data (B). Omic data was normalized using relative log expression (RLE), numbers in the cell are *R^2^* values as percents, and a star indicates a significant association by a permanova test

**Supp. Figure 3.**
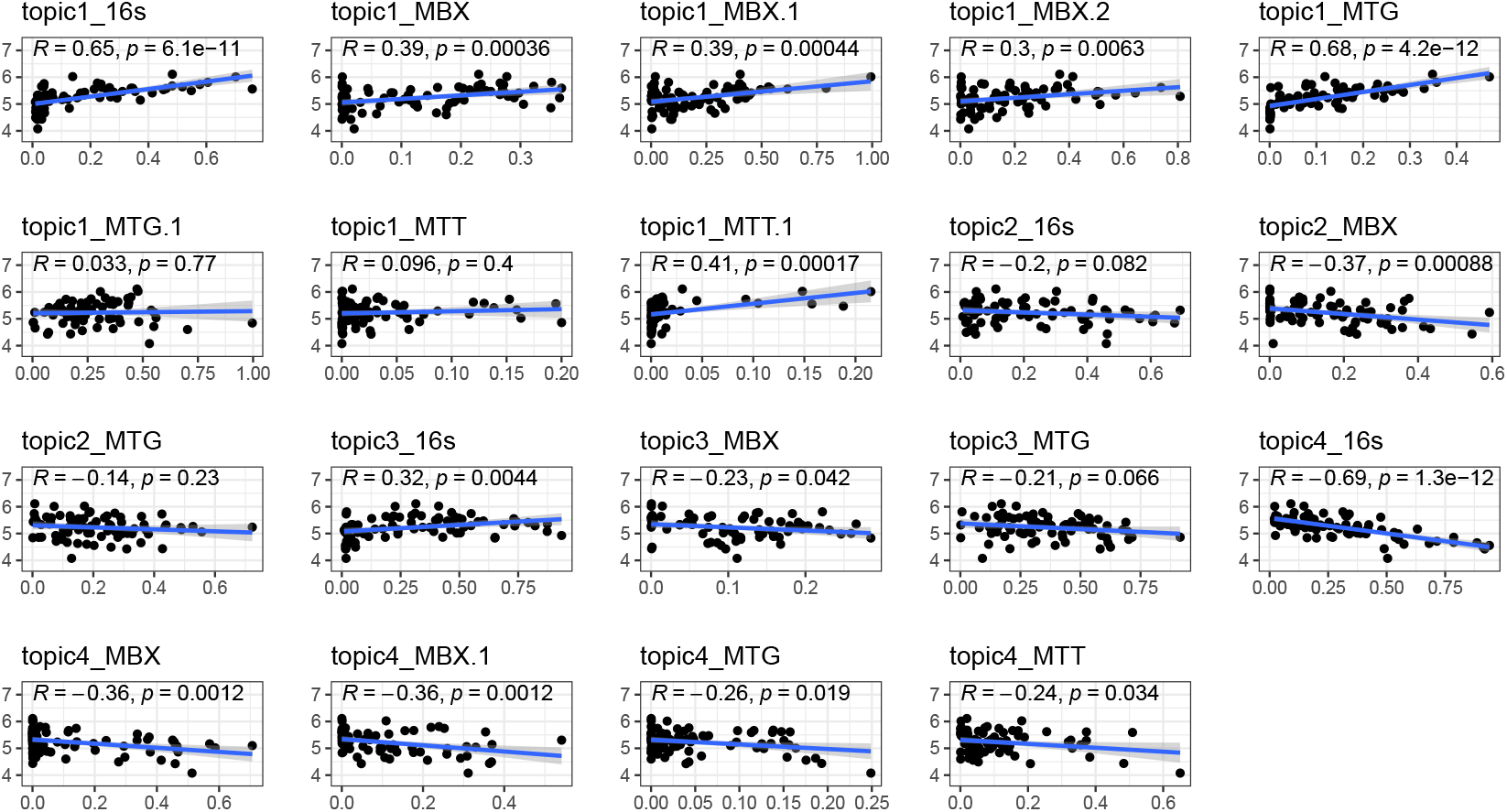
Correlation between topic values and 16S Shannon diversity. Lines and confidence intervals are linear regression fits, while R and p values are spearman correlations.

**Supp. Figure 4.**
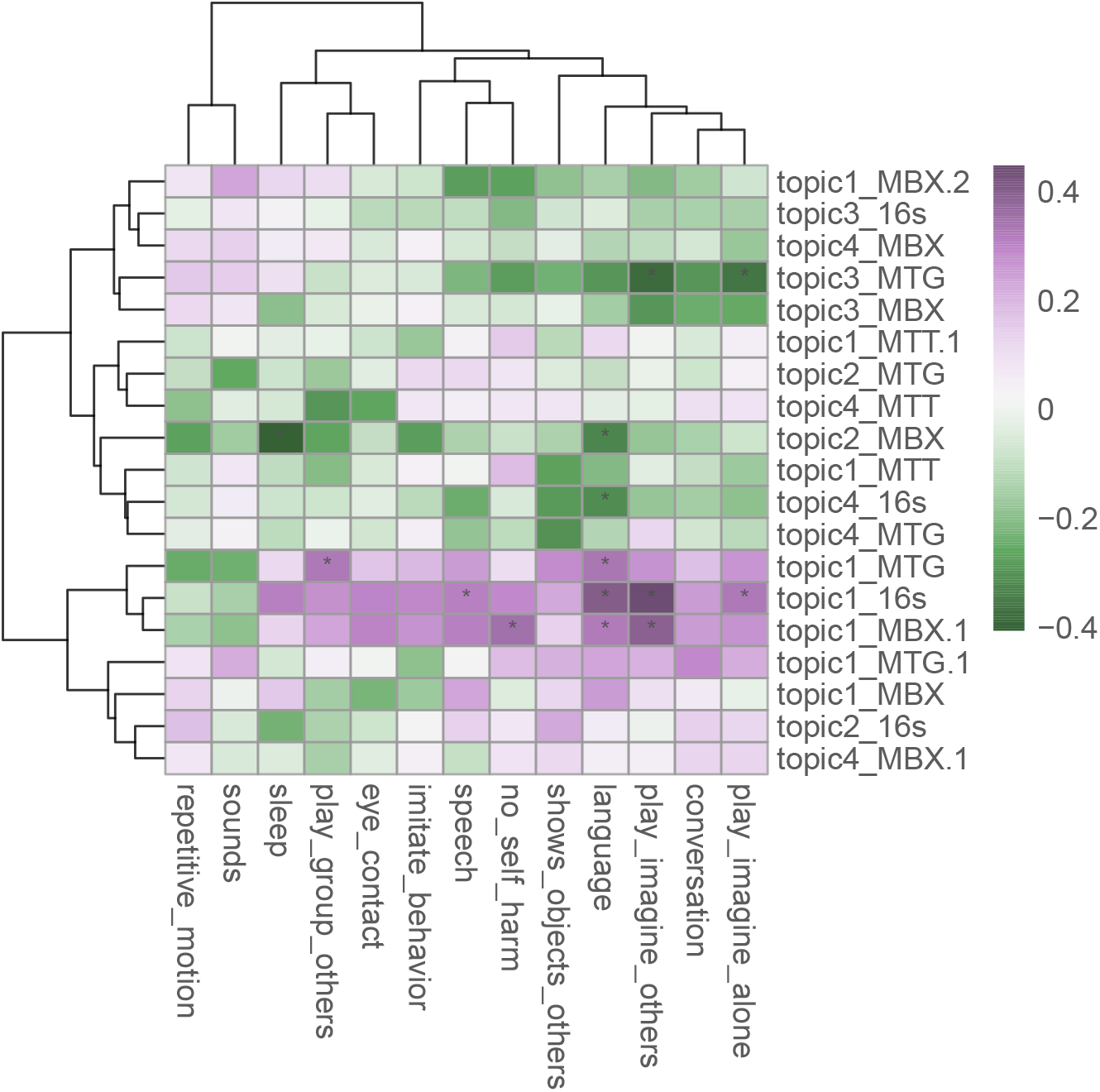
Correlations between topics and specific Autism-related behaviors, as determined by parent-completed questionnaire. Questionnaire was only completed for autistic children. Cells are colored by the spearman correlation test *R^2^* value, and a star indicates significant (p<0.05).

**Supp. Figure 5.**
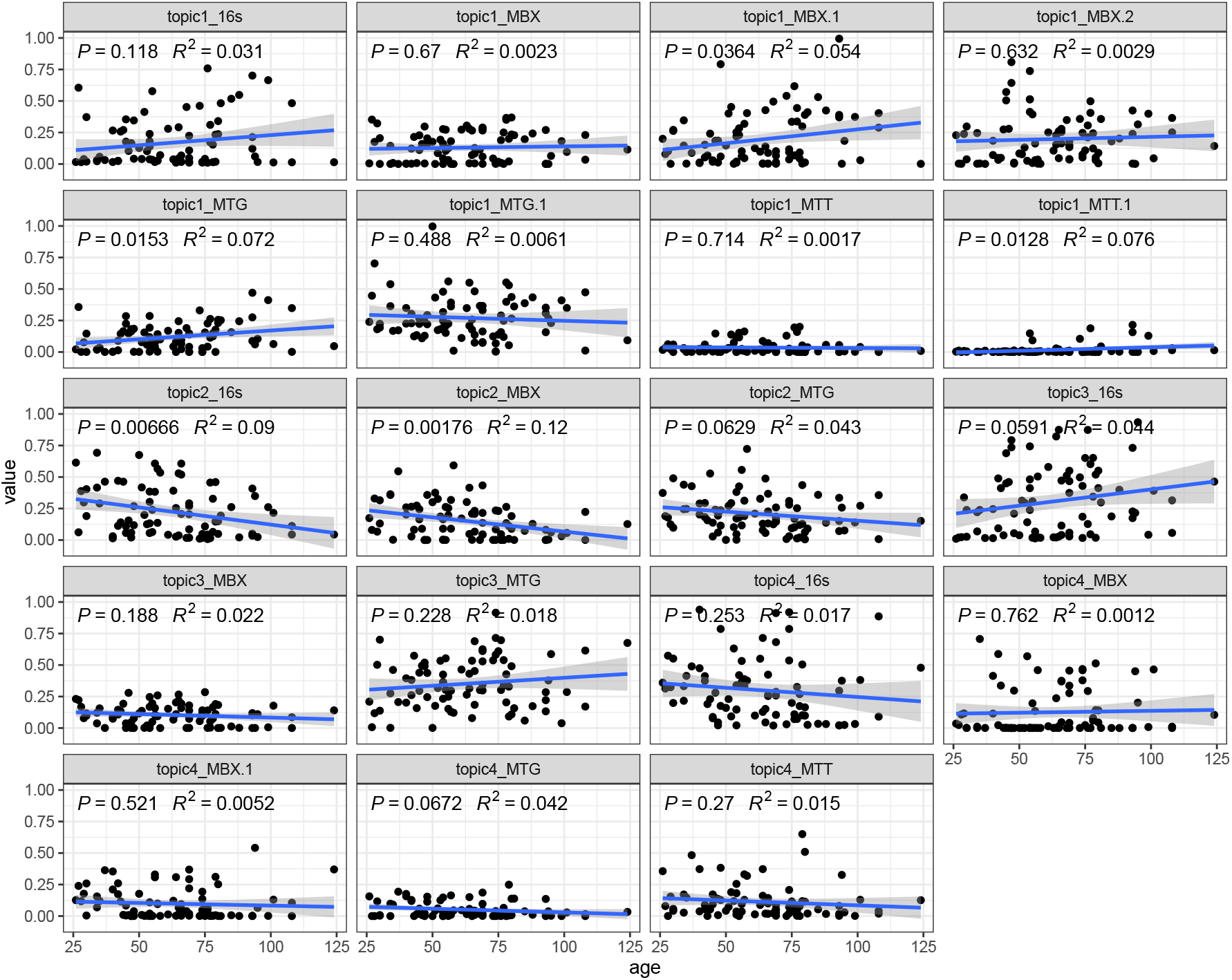
Spearman correlation between age (in months) and topic values. Topic 2 most consistently correlates inversely with age, and Topic 1 correlates with age in some omics.

**Supp. Figure 6.**
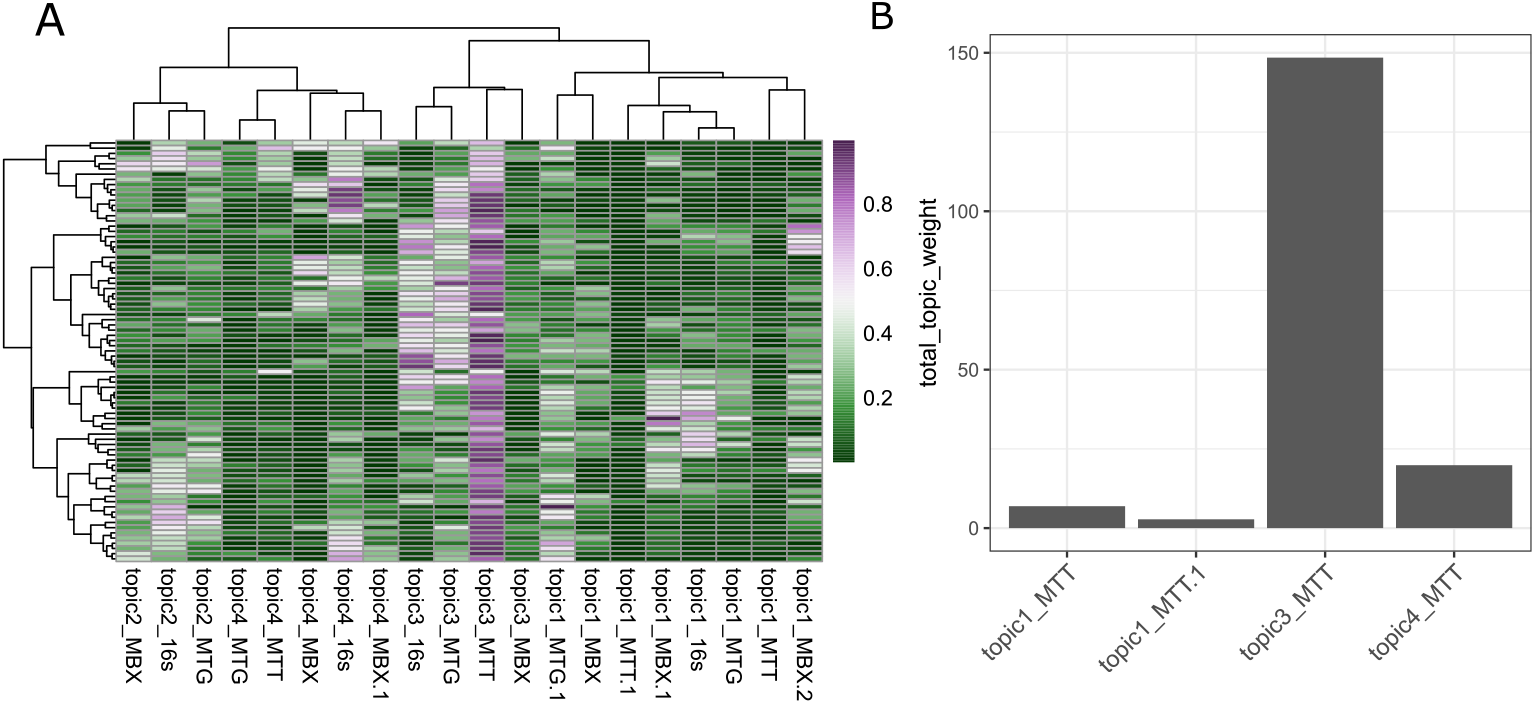
MTT topic 3 was removed due to failure to low variability across samples and excessively high topic attribution across samples. MTT topic 3 has high topic values across nearly all samples, and far exceeds all other topic value magnitudes (A). MTT topic 3 has orders of magnitude higher topic attribution across samples compared with other MTT topics (B).

**Supp. Figure 7.**
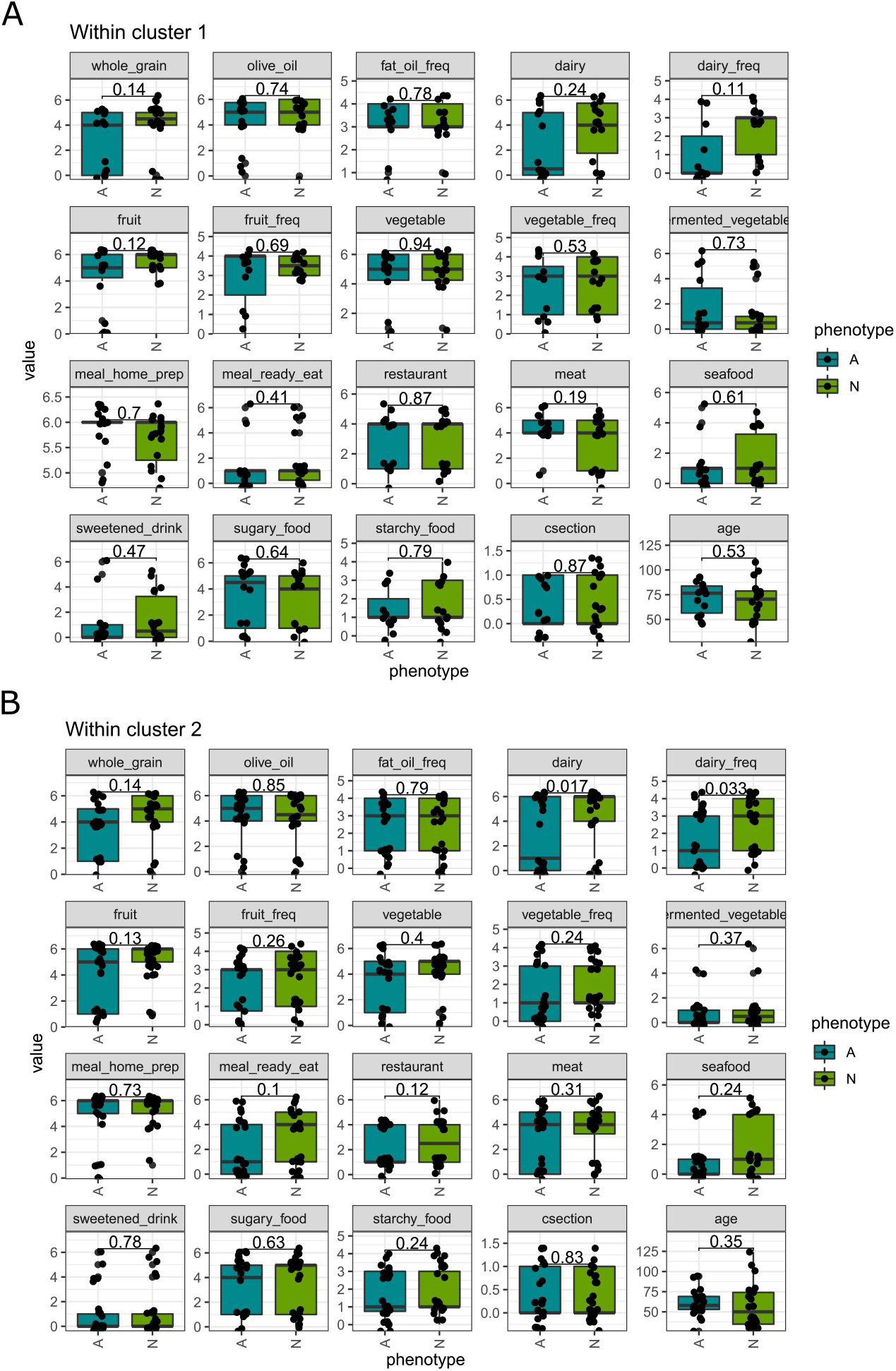
Wilcox rank sum test on dietary variable differences between Autistic and typically developing participants per sample cluster.

**Supp. Figure 8.**
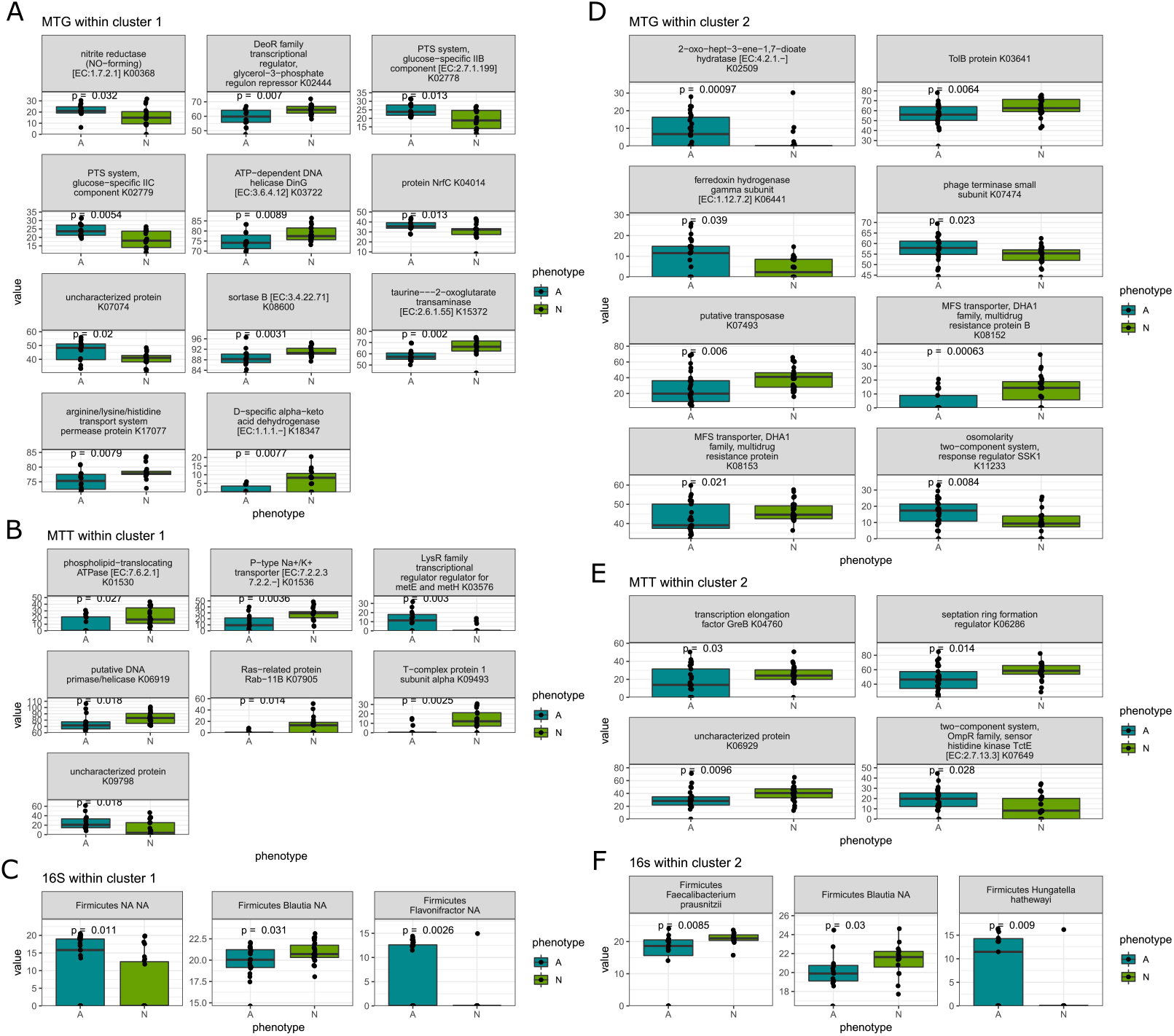
Differential analysis of MTG, MTT, and 16S features between autistic and typically developing children within sample clusters as determined by topics. Within cluster 1, MTG features (A), MTT features (B), 16S features (C). Within cluster 2, MTG features (D), MTT features (E), 16s features (F). Features are first filtered using the Boruta package, and then p values are reported using Wilcox rank sum test.

**Supp. Figure 9.**
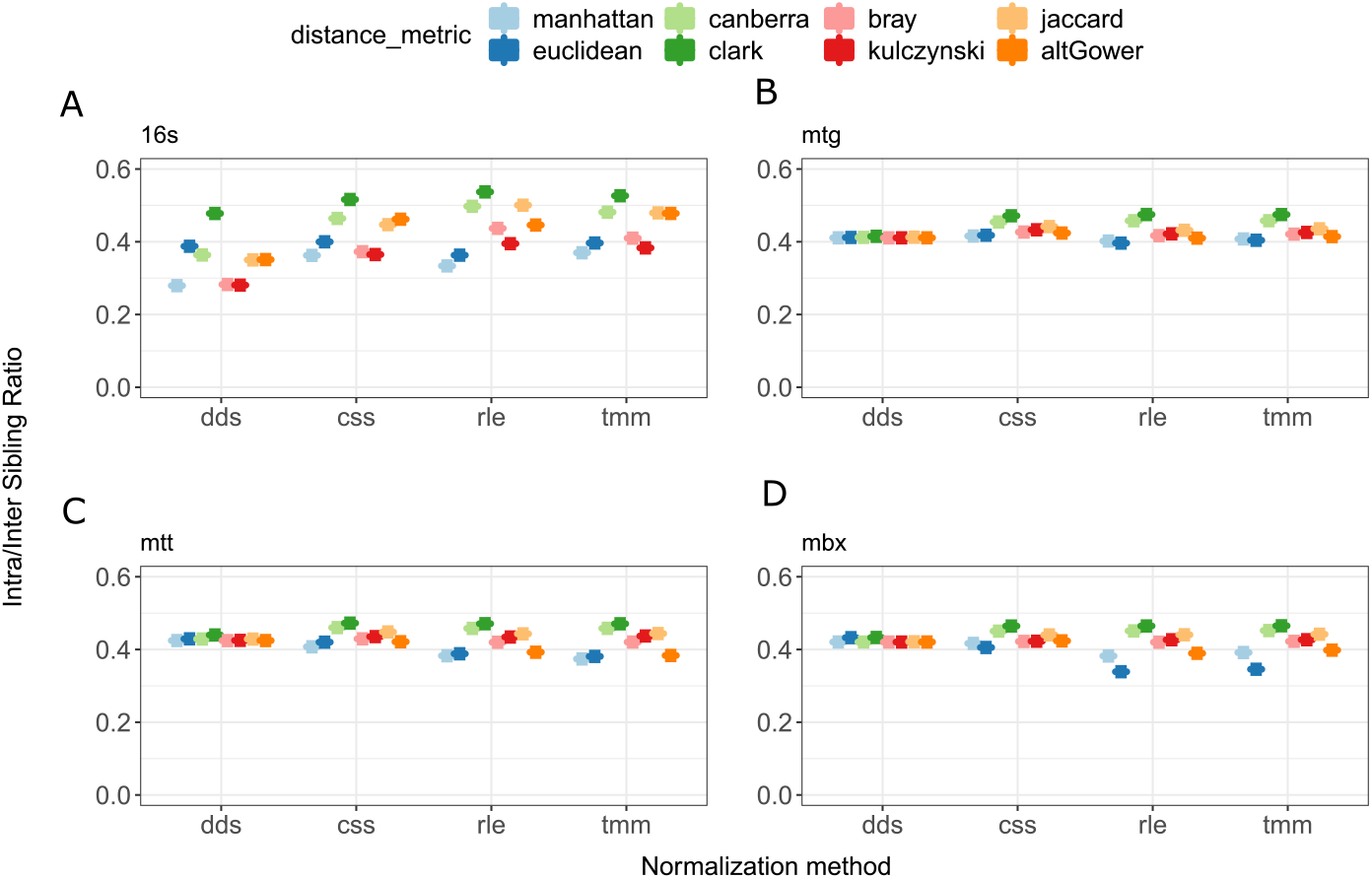
Choosing a normalization method per omic datasets. Normalization methods that minimized the distance between siblings vs. the distance between all unrelated samples there selected. When differences in normalization performance were negligible, the majority method was used. 16S DeSeq2 (A), MTG RLE (B), MTT RLE (C), MBX RLE (D)

**Supp. Figure 10.**
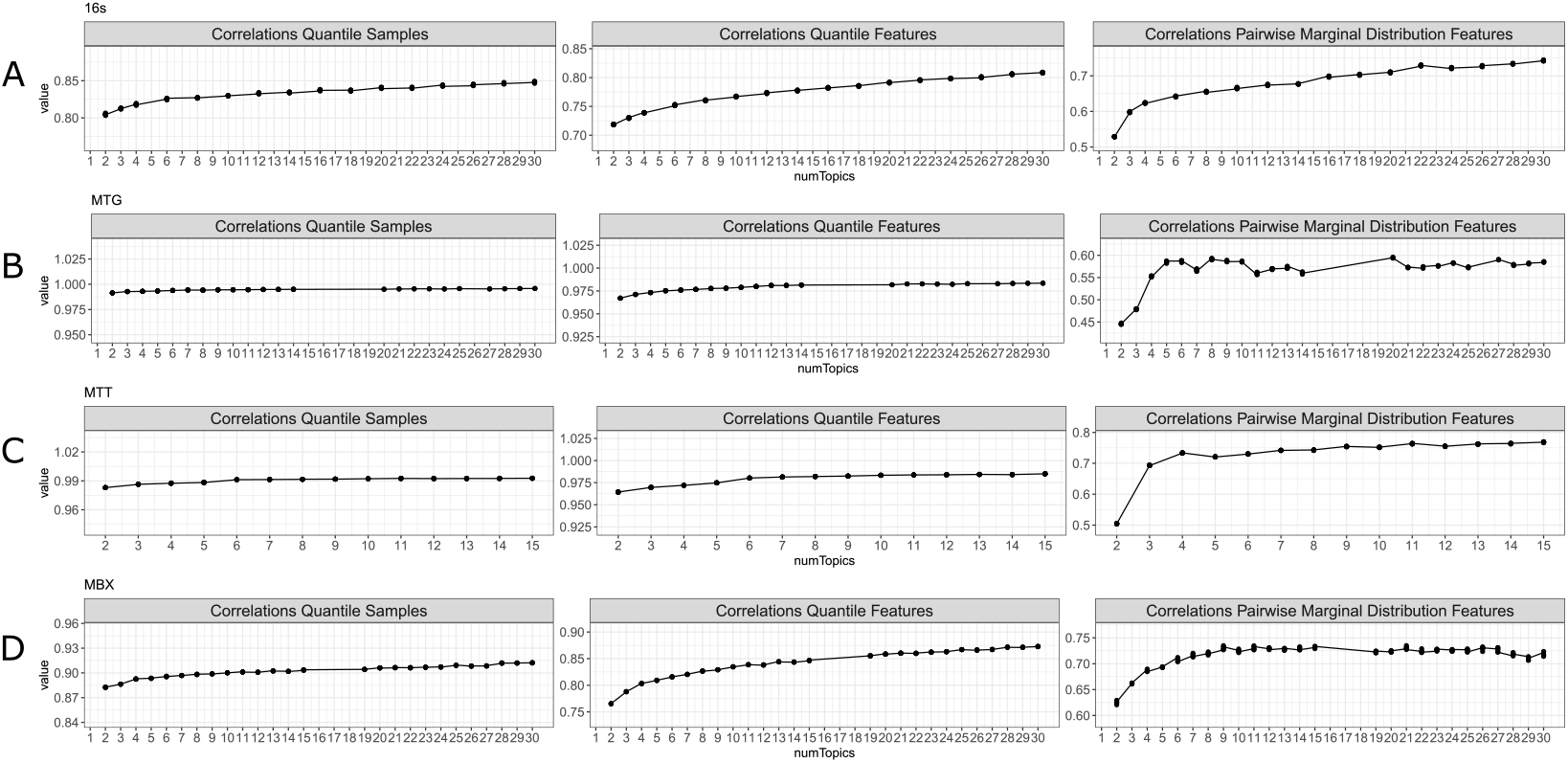
Selecting number of topics per model. To evaluate model fit, sample data was simulated using each model, and simulated and true data were compared using three metrics: correlation of sample quantiles, correlation of feature quantiles, and marginal pairwise distribution of features. Number of topics was selected using the elbow method. The final number of topics selected were 4 for 16s data (A), 5 for MTG data (B), 4 for MTT data (C), and 7 for MBX data (D).

